# Multi‑omic and spatial analysis reveals tumour‑derived paracrine signals drive suppressive macrophage polarisation via activation of the cAMP-CREB axis in glioblastoma

**DOI:** 10.64898/2026.02.19.706754

**Authors:** Samuel S. Widodo, Marija Dinevska, Stanley S. Stylli, Riccardo Dolcetti, Roberta Mazzieri, Pouya Faridi, Terry C.C. Lim Kam Sian, Stefano Mangiola, Laraib A. Ali, Sabine Vettorazzi, Jan Tuckermann, Marlene Hao, Lincon Stamp, Miguel A. Berrocal-Rubio, Alexander D. Barrow, Laura Cook, Theo Mantamadiotis

## Abstract

Tumor-associated macrophages (TAMs) are key mediators of tumor immunosuppression, yet the factors governing their polarization remain poorly understood, especially in highly immunosuppressive cancers, including cancers affecting the central nervous system. This study investigates the molecular pathways underlying TAM polarization in glioblastoma, one of the most immunosuppressive cancer types. Using a multi-omics approach integrating spatial proteomics, RNA-sequencing, and proteomic profiling of tumor cells and macrophages, we demonstrate that circulating monocytes polarize toward an immunosuppressive state when they exit tumor blood vessels, in response to glioblastoma cell-secreted cytokines. In situ and in vitro data shows that macrophage polarization is regulated via the cAMP-CREB signaling-transcription axis.

## Introduction

Glioblastoma (GBM) is one of the most aggressive cancers, with a median survival of 14.6 months from diagnosis, and the standard of care treatment includes maximal safe surgical resection and concurrent chemoradiotherapy (Stupp et al., 2005). GBM is characterized by a robust immunosuppressive tumor microenvironment (TME) largely dictated by the extensive infiltration of tumor-associated macrophages (TAMs), which includes both tissue resident microglia and bone marrow-derived macrophages (BMDM). Although both microglia and BMDM are present in the GBM TME, the consensus view is that BMDM are the major myeloid cell type responsible fueling tumor immunosuppression in GBM, given that these cells can constitute up to 50% of the cells in GBM, and that BMDM exhibit robust immune suppression while microglia exhibit weaker immune suppressive function, compared to BMDM ^1, 2,3^. Single cell sequencing of immune cells from patient GBM, distinguishing tumor core and tumor periphery, demonstrated that BMDM are the predominant infiltrating immune cell within the tumor core, whereas TMEM119^+^ microglia are largely restricted to the tumor periphery ^4^. In tumors, a highly immunosuppressive TME hinders the development of both intrinsic and immunotherapy-induced anti-tumor immune responses. Indeed, tumor immunosuppression is one of the key cancer hallmarks: avoidance of immune sensing/destruction ^5^. Modulation of TAM polarization could be accomplished through the targeted inhibition of specific cell signal transduction cascades and associated downstream transcriptional regulators ^6^.

A key characteristic of macrophages is their functional plasticity, which is dependent on stimuli within the TME, regulated via complex cell signaling and transcriptional programs ^8,9^. A transcription factor which acts as a downstream hub of multiple cytoplasmic signaling pathways is the cAMP response element binding protein (CREB) ^10^. In the context of immunity, CREB is reported to regulate inflammatory responses via activation of the cAMP and MAPK pathways ^11,12^. In macrophages, CREB can directly regulate the expression of anti-inflammatory factors, including IL-10 ^13^ and VEGFA ^14^. A recent study demonstrated that pharmacological inhibition of protein kinase A (PKA), a key cAMP pathway effector protein and upstream activator of CREB, reduced the expression of several anti-inflammatory factors in breast cancer TAMs, including IL10, VEGFA and ARG1, and prolonged survival in mice ^15^.

In this study we characterized CREB activation in BMDM TAMs, in the GBM TME using spatially resolved multiplex immunohistochemistry to distinguish M2-type immunosuppressive TAMs from M1-like inflammatory TAMs and determined the CREB activation status of these cells. Spatial analysis showed that circulating BMDM TAMs are triggered to polarize toward an immunosuppressive state once they exit the circulation and encounter neoplastic cells in the tumor parenchyma. The majority of BMDM in the tumor parenchyma were phospho-CREB-positive, while BMDM located within tumor blood vessels and perivascular niches were phospho-CREB-negative, suggesting that CREB regulates a shift from an M1-like state to an M2-state in response to factors within the TME. Spatial tissue analysis identifying cell signaling activation in BMDM combined with in vitro experiments support the view that BMDM M2-like polarization by GBM-cell secreted factors is regulated by cAMP signaling but not by PI3K and MAPK signaling.

To identify GBM cell-secreted factors which regulate macrophage polarization, human leukemia monocytic THP-1 cells and peripheral blood mononuclear cells (PBMC) were used to generate monocyte-derived macrophages (MDM). Exposure of MDM to GBM conditioned medium was sufficient to polarize macrophages toward an M2-like immunosuppressive state. GBM conditioned medium-treated MDM cells incubated with and without CREB inhibitor and analyzed for mRNA and protein expression suggest that CREB regulates M2-like TAM polarization, by regulating the expression and secretion of immunosuppressive cytokines and chemokines.

Overall, this is the first study which defines tumor niche-specific cell signaling and transcriptional changes occurring in macrophages, as they transition from a tumor-naïve state to immunosuppressive tumor associated macrophages. We show that tumor secreted cytokines are sufficient for triggering M2-type macrophage polarization, via the cAMP-CREB signaling-transactivation axis. Together with previous data demonstrating that CREB regulates glioma malignancy in a high-grade glioma mouse model, our data suggests that inhibiting CREB has the potential to target both neoplastic cells and reprogram the tumor microenvironment.

## Methods

### Brain tumor tissue

Human brain tumors and non-tumor formalin fixed paraffin-embedded (FFPE) tissue sections were obtained from the Department of Anatomical Pathology, Royal Melbourne Hospital and from US Biomax (now TissueArray.Com LLC). Human ethics approval for the use of these specimens was covered by project application 1853511 and was approved by the Medicine and Dentistry Human Ethics Sub-committee, University of Melbourne. Fifty-nine patient tissues were used: non-tumor (tumor adjacent normal cerebrum tissue), n=4; grade 2, n=20; grade 3, n=7; GBM, n=28. All tissue was assessed by an anatomical pathologist. IDH mutation status of all glioma/GBM tissue was determined, as previously described ^16^.

### Human Ethics

Use of human tissue samples and data received ethics approval from the University of Melbourne Human Research Ethics Committee. Use of human brain tumor tissue, and data derived from these, is covered by ethics project #28992. The use of healthy donor blood and PBMC is covered by ethics projects #49519, #39869 and #61740.

### Multiplex immunohistochemistry and image acquisition

Multiplex immunohistochemistry was performed using an Opal 7-colour IHC kit (Akoya Biosciences) on a Bond-RX auto-stainer (Leica). 4μm thick FFPE tissue sections were deparaffinized in xylene and rehydrated in a serial dilution of ethanol (100%, 100%, 80%, 30%). Heat induced-antigen retrieval was then performed in either citrate buffer pH6 or Tris-EDTA buffer pH9 for 15 minutes at 125°C. Sections were incubated in 3% hydrogen peroxide to block endogenous peroxidase activity and primary antibody diluted in antibody diluent solution (Akoya Biosciences) was added. A CREB macrophage antibody panel was used to investigate the expression of pCREB in subsets of macrophages and their localization in brain tumor samples. The antibody panel is shown in Table S1. The primary antibody was incubated for 1h at room temperature, followed by incubation of Opal polymer horseradish peroxidase Ms+Rb secondary antibody for 30 min. Opal fluorophore, diluted at 1:150 in 1x Plus Amplification Diluent (Akoya Biosciences), was applied for 30 minutes at room temperature. Serial multiplexing was performed by repeating the heat induced-antigen retrieval step, followed by application of different primary antibodies, Opal polymer, and Opal fluorophore. Spectral DAPI (1 drop in 500μL PBST) was added and incubated for 5 minutes, and sections were mounted in CitiFluor CFPVOH Solution plus antifade (CitiFluor, Australia). Images were acquired using a Vectra 3.0.5 multi-spectral imaging platform (Perkin Elmer, USA), at 40x magnification.

### Image analysis

Whole slide multi-spectral images acquired using the Vectra platform were deconvoluted using inForm v2.4.8 software (Akoya Biosciences) and image tiles were then fused and analyzed using HALO software (Indica Labs). Multiplex immunohistochemistry image analysis was performed using the Highplex FL 3.0 algorithm module in HALO. Signal thresholding of each biomarker fluorophore channel was also performed to define true positive staining, perform cell phenotyping and cell counting, in annotated tissue regions. For spatial analysis, serial Masson’s Trichrome stained images were overlaid with multiplex immunohistochemistry images to define the perivascular collagen-rich regions. Collagen-rich regions, identified as the green areas in the Masson’s Trichrome stained tissue, were defined as ECM-rich regions, and cellular regions lacking ECM were defined as neoplastic cell (NC)-rich regions. Cell preferential distribution analysis was performed by enumerating cell types in both perivascular ECM-rich and NC-rich regions.

### Cell line culture

Human GBM cell lines (U-87MG, LN-229, LN-18, U-118MG, T98G, A172) obtained from the ATCC, and patient-derived GBM cells (MU41) ^17^ were maintained in Dulbecco’s Modified Eagle’s Medium (DMEM) (Thermo Fisher Scientific) with 10% fetal calf serum (FCS) (Bovogen, Australia, #SFBS-F) and penicillin-streptomycin-amphotericin B (Gibco Anti-Anti, Thermo Fisher Scientific). Human monocytic THP-1 cells were cultured in RPMI (Thermo Fisher Scientific) with 10% FCS and penicillin-streptomycin-amphotericin B. Tumor conditioned medium was produced by seeding GBM cell lines in Opti-MEM reduced-serum medium (Thermo Fisher Scientific) at 60% confluency, following 72 h of culture.

### THP-1 cell macrophage differentiation

THP-1 cells were seeded at 1×10^5^ cells in a 6-well plate in RPMI + 10% FCS. Cells were treated with 100 ng/mL phorbol 12-myristate 13-acetate (PMA) for 24 h to induce differentiation to macrophages. Following treatment, adherent cells were washed with PBS buffer and added serum-free RPMI for a 48-hour resting period. After resting, media were replaced with 50% of GBM conditioned medium in RPMI. Cells were then collected for analysis after 30 minutes. Macrophages were exposed to GBM cell conditioned medium for 6 h for RNA analysis and 24h for protein analysis.

### Generation of monocyte-derived macrophages

Buffy coat samples were obtained from healthy donors aged between 25 and 40 years old. For PBMC isolation, blood was mixed with cold PBS (Media Preparation Unit, University of Melbourne) at 1:1 ratio. Approximately 12.5mL Lymphoprep (Stemcell Technologies) was added to the bottom of a 50mL tube and 25mL blood + PBS mixture was then slowly layered on top. Tubes were centrifuged, with slow acceleration and deceleration, at 600g for 20 min at room temperature.

After centrifugation, the PBMCs were then transferred into a new tube using a transfer pipette and mixed with PBS (Media Preparation Unit, University of Melbourne). After the PBMCs were collected, the cells were suspended in 50ml PBS and centrifuged at 500g for 7 minutes at 4°C. Red blood cells (RBCs) were removed by lysis using RBC lysis buffer (1500mM NH_4_Cl, 1mM KHCO_3_, 10mM EDTA). Purified PBMCs were resuspended in 50ml PBS and counted using a hemocytometer.

CD14^+^ monocytes were enriched from isolated PBMCs using EasySep Human CD14 Positive Selection Kit II (Stemcell Technologies). Monocytes were then cultured in complete RPMI supplemented with 20ng/ml recombinant human CSF-1 (Peprotech) for 6 days. Cell culture media was replaced every 3 days. On day 6, the medium was replaced with serum-free RPMI and the cells allowed to rest for 48 h before further experimental treatment.

### Western blotting

Whole cell lysates were prepared by lysis in PRO-PREP Protein Extraction Solution (Intron Biotechnology) supplemented with 1x Halt Protease and Phosphatase Inhibitor Cocktail (Thermo Fisher Scientific). Lysates were then cleared from cell debris by centrifugation at 12,000 rpm for 10min at 4°C. Concentration of the cleared lysates was measured using Pierce BSA protein assay kit (Thermo Fisher Scientific) as described by the manufacturer.

Protein samples were prepared in NuPAGE LDS Sample Buffer (Invitrogen) and NuPAGE Sample Reducing Agent (Invitrogen). Samples were heated at 95°C for 5 minutes, pulse-spun and 20μg total protein loaded onto 10% SDS PAGE gels. Gels were run at 90V for 1.5h, in SDS running buffer. Proteins were transferred onto nitrocellulose membranes (Amersham Pharmacia). Membranes with transferred proteins were then blocked using 5% skim milk in Phosphate Buffer Saline supplemented with 0.1% Tween-20 (PBST). Primary antibodies (1:10,000) were then added and incubated overnight at 4°C. On the following day, membranes were washed in PBST, followed by incubation of secondary antibodies conjugated with HRP for 1 hour at room temperature. Membranes were then washed in PBST at room temperature and proteins were detected and visualized using ECL VisULite (R&D Systems). Uncropped immunoblot images are provided as source data.

### RNA sequencing and analysis

Cells were stained with antibodies targeting biomarkers shown in Table S1. Live viable cells were then sorted using CytoFLEX SRT (Beckman Coulter) and collected in PBS supplemented with 2% FCS. Cells were then spun, and pellets were stored in -80°C. RNA purity was determined by measuring the ratio of absorbance values at 230, 260 and 280nm. Purified RNA with a A260/A280 ratio above two was used for sequencing. RNA integrity was examined using an Agilent Bionanalyzer or PerkinElmer LabChip GX. RNA was then sequenced using NovaSeq X Plus, at a depth of >20M reads.

RNA sequence quality was first determined using FastQC (Andrews, 2010). Sequences were then aligned to the Genome Reference Consortium Human Build 38 patch release 14, obtained from the Genome Reference Consortium (NCBI RefSeq assembly GCF_000001405.40). Count data was then generated using the R package featureCounts (Liao et al., 2014) and further downstream analyses were performed using the *tidyomics* software ecosystem (Hutchison et al., 2024), including principal component and differential gene expression analysis.

### Cytokine analysis

To analyze the secreted factors produced by GBM cells and GBM-stimulated macrophages, protein analysis using the Olink Proximity Extension Assay system and an Olink system Target 96 Inflammation panel was performed on conditioned media from GBM cells and GBM cell conditioned medium-stimulated macrophages. These experiments were performed at Monash Proteomics and Metabolomics Platform, Monash University. The amount of each factor measured was determined and expressed as Normalized Protein eXpression (NPX).

Data analysis and visualization was performed in RStudio using the R package, OlinkAnalyze (https://cran.r-project.org/package=OlinkAnalyze).

Cytokine levels in cell supernatants were also analyzed using the BioLegend LEGENDplex (San Diego, USA) cytometric bead array (CBA) Human inflammation panel 1 kit (13-plex #740809). The analytes measured were IL-1β, interferon (IFN)-α2, IFN-γ IL-8, TNF-α, MCP-1, IL-6, IL-8, IL-10, IL-12p70, IL-17A, IL-18, IL-23, and IL-33. The level of soluble cytokines was determined using a 4-laser CytoFLEX S cytometer (Beckman Coulter, USA).

### Protein size fractionation

To examine the content of GBM cell conditioned medium (CM), MU41-CM was molecular size-fractionated using Amicon ultracentrifuge filters (Merck) with molecular weight cut-offs of 100kDa, 30kDa and 10kDa. Filtered CM fractions were then added to THP-1 macrophages. MU41-CM was also treated with Proteinase K to determine whether macrophage-active CM components were proteins. Unfiltered MU41-CM and CM-free basal media were used as positive and negative controls, respectively.

## Results

### Shift toward M2-type macrophage polarization during disease progression and migration into the GBM tumor parenchyma

Using histological staining and multiplex immunohistochemistry to label cells co-expressing CD68, CD163, CD206, TMEM119 and HLA-DR (Fig 1A,B,C), we examined MDM (CD68^+^

**Figure 1.**
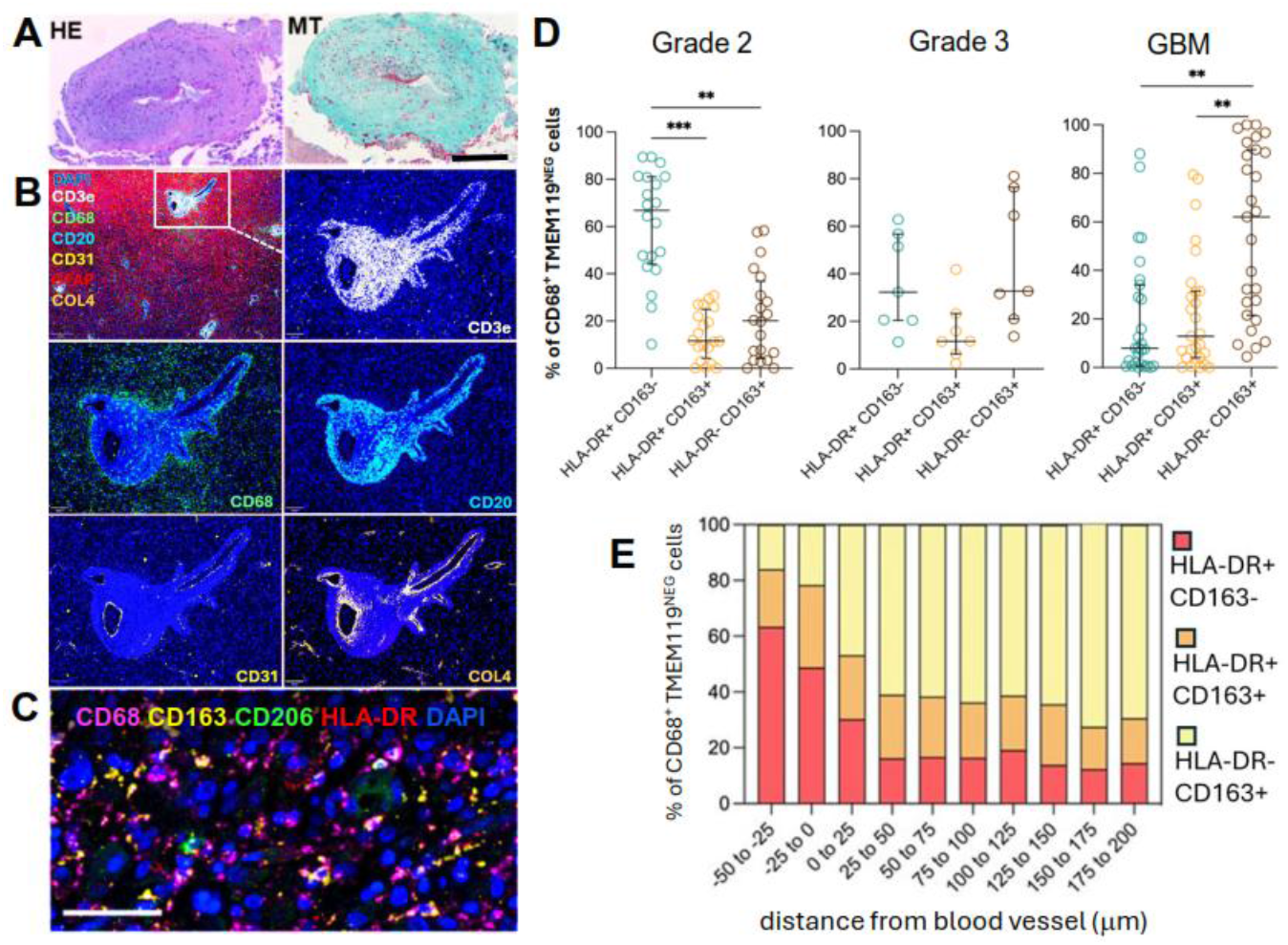
Immune cell infiltration and TAM polarization in GBM. **(A)** HE and Masson’s trichrome stained GBM tissue showing blood vessel and perivascular niche with ECM/collagen (green). Scale bar is 100μm. **(B)** multiplex immunofluorescence image of GBM tissue stained using antibodies recognizing CD68, HLA-DR and CD163. Scale bar is 50μm. **(C)** Multiplex immunofluorescence image of GBM tissue stained for myeloid biomarkers CD68, HLA-DR and CD163 and CD2026. **(D)** CD68^+^ TAM subsets which are HLA-DR^+^, HLA-DR^+^ CD163^+^ double positive and CD163^+^ in grade 2, 3 astrocytoma and GBM tissue. **(E)** Infiltration analysis showing TAM subset density up to 200mm from blood vessels in GBM tissue. Data is presented as infiltration distance from the outer border of CD31^+^ vascular cells (-50 to 0mm), into NC-rich regions (0 to 200mm).

TMEM119^NEG^ cells) status in grade 2 (n=20), grade 3 (n=7) astrocytoma tissues and GBM (n=28), respectively. We observed a predominance of HLA-DR^+^ CD163^NEG^ TAMs in grade 2 tumors (67% of TAMs), and a shift toward cells expressing CD163 and no HLA-DR (HLA-DR^NEG^ CD163^+^) in grade 3 astrocytoma and GBM, where 32% (grade 3 astrocytoma) and 63% (GBM) of TAMs were HLA-DR^NEG^ CD163^+^ (Fig 1D). Cells co-expressing HLA-DR and CD163 (HLA-DR^+^ CD163^+^) comprised between 10 and 15% of all TAMs, across all tumor grades.

To determine if macrophage polarization is regulated by the TME, we performed spatial cell infiltration analysis by measuring cell density around blood vessels, in 28 GBM patient tissues (see Materials and Methods). Within tumor blood vessels, located between -50μm and 20μm, between 50% and 60% of TAMs, identified as CD68^+^ TMEM119^NEG^ cells, were HLA-DR^+^ CD163^NEG^ (Fig 1E). Infiltration analysis of CD68^+^ TMEM119^NEG^ cells, showed that TAM polarization shifted toward an immunosuppressive, M2-like state in the tumor parenchyma, where between 60% and 70% of TAMs were HLA-DR^NEG^ CD163^+^. CD68^+^ cells co-expressing HLA-DR and CD163 (HLA-DR^+^ CD163^+^) comprised between 20% and 30% of TAMs within blood vessels, and between 15% and 20%, at distances at and beyond 25μm of blood vessels, within the tumor parenchyma.

### Phospho-CREB^+^ macrophages in the GBM tumor parenchyma co-express CD163 and CD206 but not HLA-DR

We previously reported that the kinase inducible transcription factor CREB exhibits a grade-dependent activation in both neoplastic cells and in infiltrating macrophages ^18^. To investigate whether CREB expression and activation correlates with macrophage polarization, we measured the proportion of bone marrow-derived TAMs expressing phosphorylated CREB (pCREB), HLA-DR and CD163. Immunofluorescence staining showed that the proportion of pCREB^+^ TAMs was highest in GBM, compared to IDH mutant grade 4 astrocytoma, and lower grade astrocytoma (Fig 2A,B). Spatial infiltration analysis showed that most pCREB^+^

**Figure 2.**
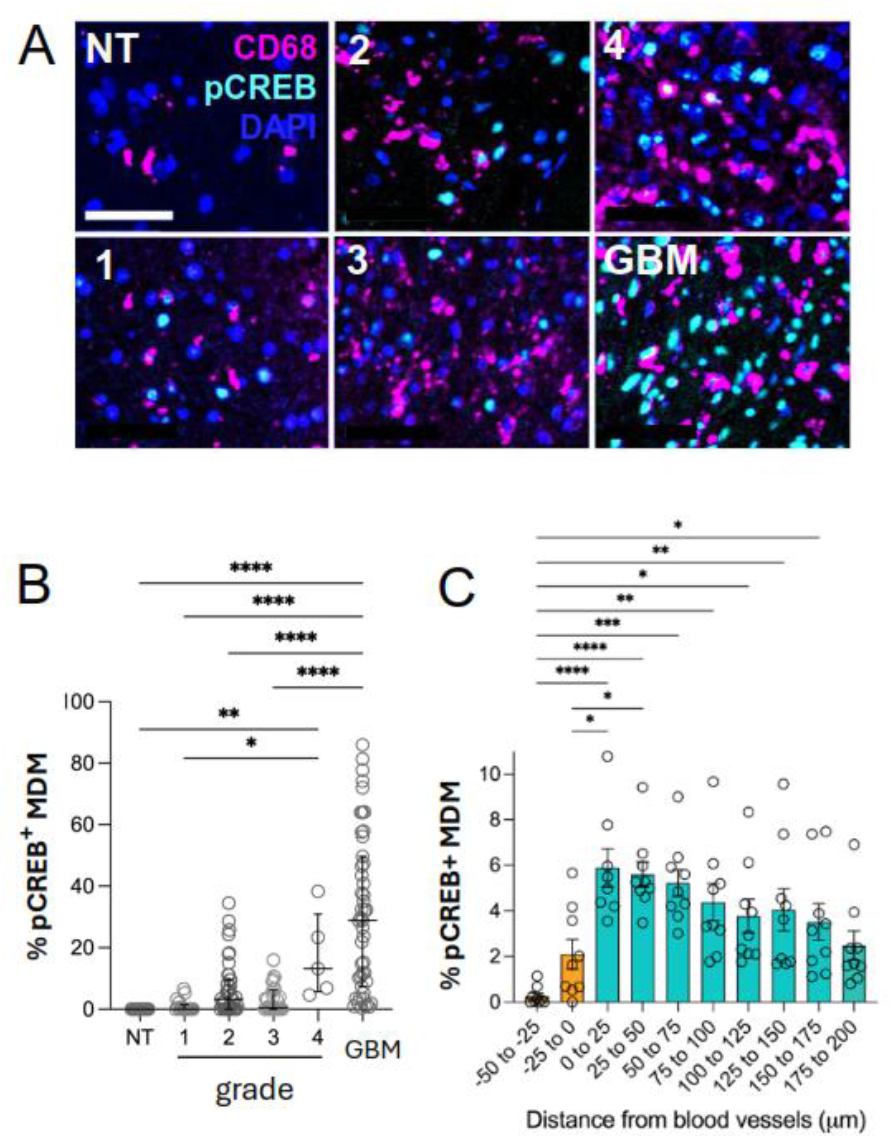
CREB activation in monocyte-derived macrophages in the GBM tumor microenvironment. **(A)** Multiplex IHC was used to identify the presence of pCREB^+^CD68^+^ MDM cells in non-tumour (NT), grades 1-4 astrocytoma and GBM tissue. Scale bar is 50mm. **(B)** pCREB^+^ MDM count from mIHC in (NT) (n=14), grade 1 (n=20), grade 2 (n=42), grade 3 (n=31) and IDH-mutant grade 4 (n=5) astrocytoma and IDH-WT GBM (n=57). **(C)** pCREB^+^ MDM infiltration/localisation in GBM (n=9) was analysed by spatial analysis and is shown as the proportion of pCREB^+^ MDM cells (Y-axis) and infiltration distance in mm (X-axis). 0mm is the outer border of a blood vessel; is -50 to 0mm cover the intra-vascular localization; positive distance values (0-200mm) represent infiltration into the tumor parenchyma.

TAMs were localized within the tumor parenchyma, and fewer than 2% of TAMs within blood vessels, located between -50μm and 20μm, expressed pCREB (Fig 2C).

To investigate the CREB activation state in TAM polarization in the GBM TME, we characterized the pCREB^+^ TAM cell population by staining GBM tissue using the myeloid/macrophage biomarkers, CD68, CD163, CD206 and HLA-DR (Fig 3A).

**Figure 3.**
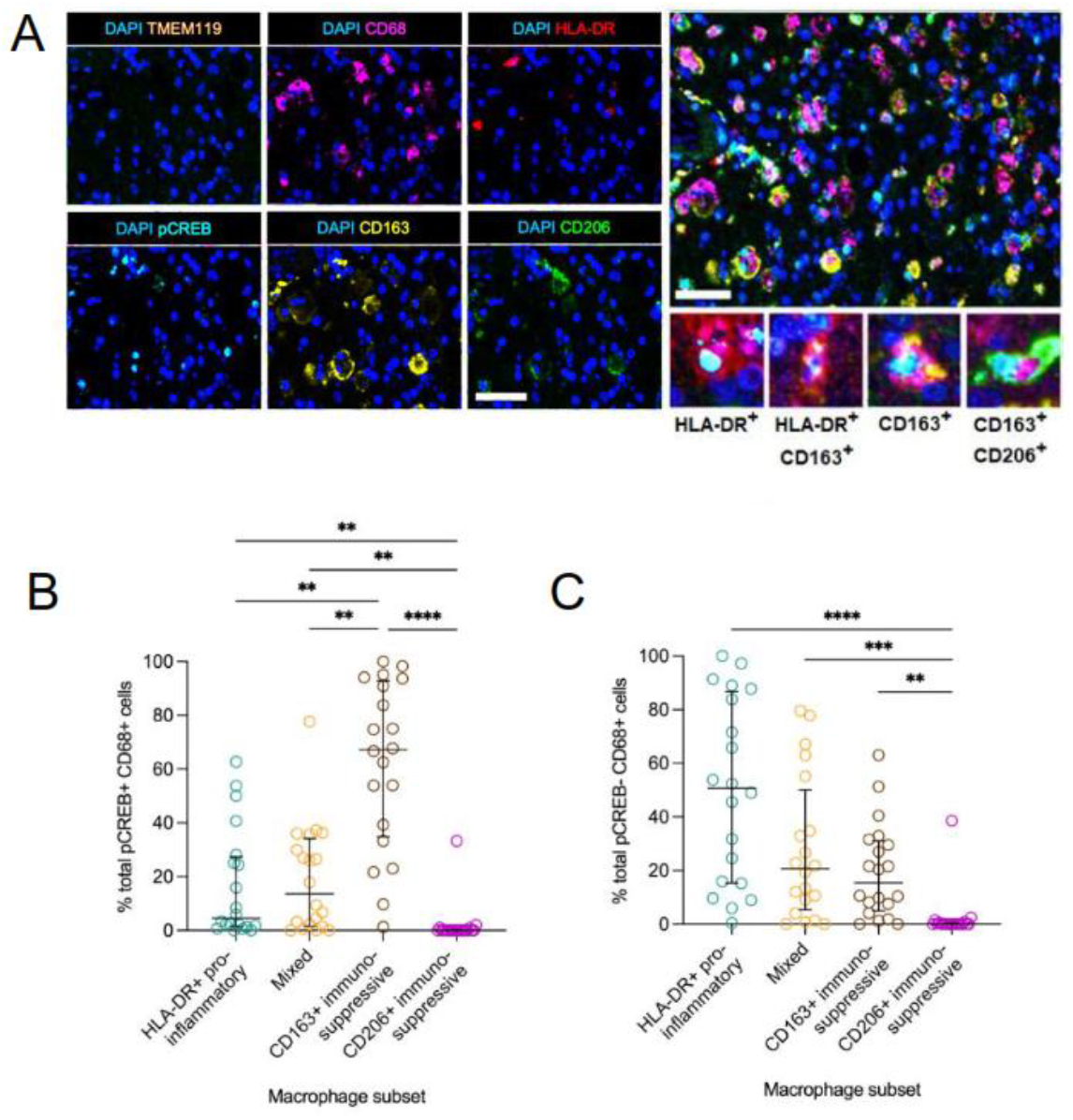
pCREB+ TAMs are immunosuppressive. **(A)** Representative mIHC images of a GBM tissue stained with antibodies labelling pCREB, CD68, CD163, CD206, HLA-DR and TMEM119. Scale bars are 50mm. **(B, C)** Proportion of macrophage subsets relative to the total pCREB+ and pCREB-(c) TAMs, in IDH-WT GBM (n=20). Error bars are median ± interquartile range. Statistical significance was determined using Kruskal-Wallis test, followed by Dunn’s test for multiple comparisons (a=0.05), and is represented by *(p<0.05), **(p<0.01), ***(p<0.001) and ****(p<0.0001).

Approximately 70% of pCREB^+^ TAMs expressed CD163 (Fig 3B), whereas most pCREB^NEG^ cells co-expressed HLA-DR (Fig 3C), suggesting that GBM-secreted factors stimulate CREB activation in TAMs.

### CREB activation in macrophages is stimulated by factors secreted by GBM cells and is regulated by cAMP signaling

To define the mechanistic role of CREB activation in tumor macrophage polarization, we used PMA-differentiated THP-1 macrophages stimulated with conditioned media from a range of patient-derived and human GBM cell lines (Fig 4A). Conditioned media from most cell lines stimulated pCREB expression and increased CREB transactivation using a luciferase CREB reporter plasmid (Fig 4B). Conditioned media from LN229, MU41 and LN18 cells exhibited the most robust CREB activation, with the weakest activation exhibited by conditioned media from U87MG and A172 cells. The differences in the ability of GBM cell conditioned media to enhance CREB phosphorylation correlated with functional CREB transactivation potential in THP-1 cells expressing a CREB luciferase reporter gene (Fig 4B). THP-1 macrophages simulated with GBM conditioned medium and co-incubated with the CREB small molecule inhibitor, 666-15 ^19^, exhibited lower expression of M2-like macrophage biomarker, CD206, compared to THP-1 macrophages, not exposed to the CREB inhibitor (Fig 4C).

**Figure 4.**
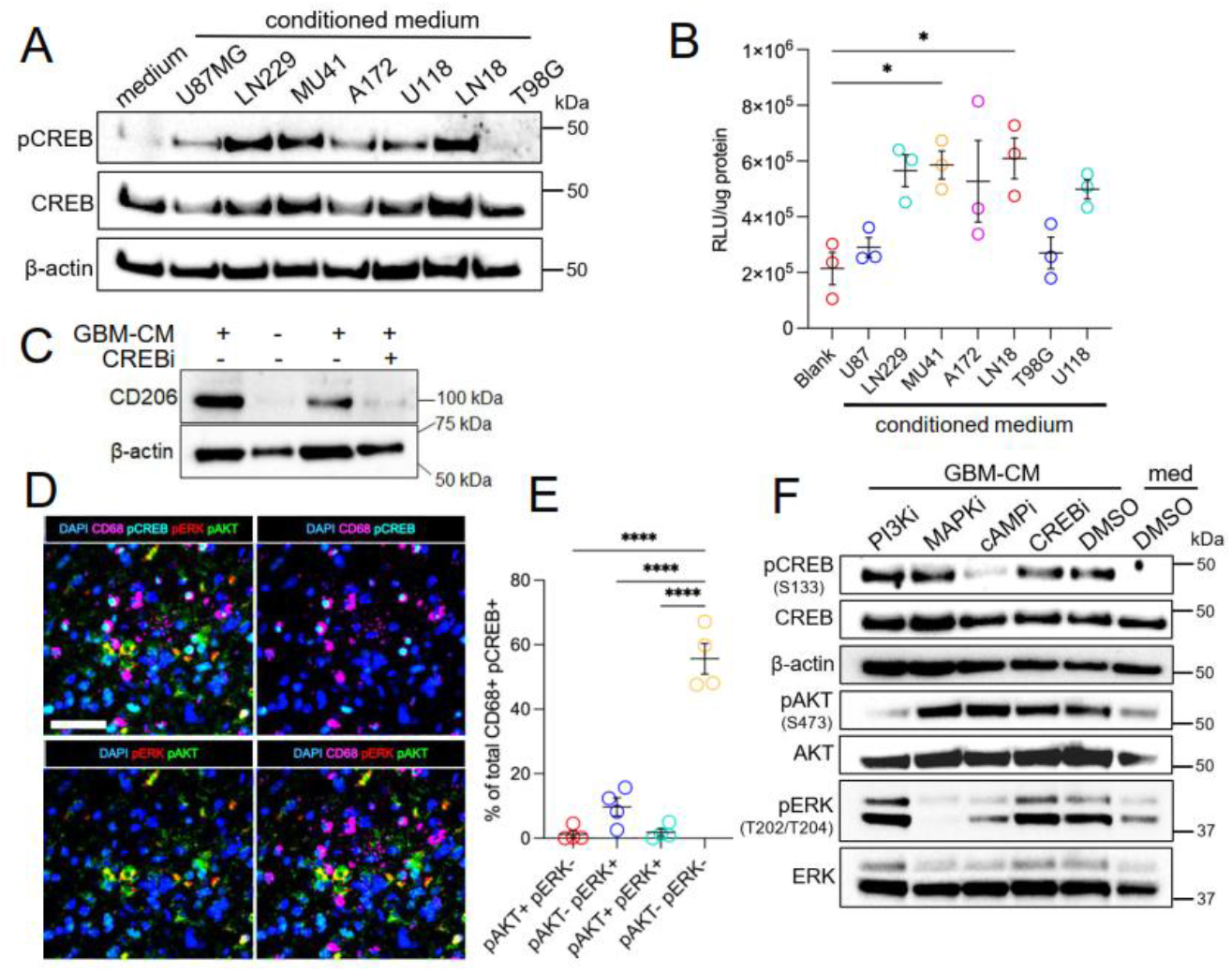
CREB transactivation by GBM secreted factors in macrophages is mediated by the cAMP signal transduction pathway. **(A)** Western blot of THP-1 differentiated macrophages stimulated for 30 min, with conditioned media (CM) from a panel of GBM cell lines and probed using antibodies for, total CREB, phospho-CREB (pCREB) and b-actin. Data shown is representative of three independent experiments. **(B)** CREB transactivation activity in THP-1 macrophages transfected with a CREB luciferase reporter plasmid after stimulation with MU41 GBM cell conditioned medium (CM). CREB activity is presented as Relative Light Units (RLU) per mg of protein lysate. Statistical significance was determined using one-way ANOVA, followed by Tukey’s test for multiple comparisons (a=0.05). *(p<0.05). Error bars are median ± interquartile range. **(C)** Expression of CD2026 in THP-1 differentiated macrophages incubated with or without GBM cell (MU41) conditioned medium (CM) in the presence or absence of the CREB inhibitor (CREBi), 666-15 (1mM). **(D)** Multiplex IHC image showing phospho-AKT (pAKT), phospho-ERK1/2 (pERK) and pCREB expression in BMDM-TAMs in GBM tissue. Scale bar is 50mm. **(E)** Percentage of CD68^+^ pCREB^+^ MDM co-expressing pAKT and pERK1/2 in GBM tissue (n=3). Error bars are S.E.M. **(F)** Cell lysates from THP-1 differentiated macrophages were probed for expression of pCREB, CREB, b-actin, pAKT, AKT, pERK and ERK following a 30-minute stimulation with GBM-CM, in combination with either a PI3K inhibitor (PI3Ki) BKM-120 (2 mM), a MAPK inhibitor (MAPKi) U0126 (10 mM), a cAMP inhibitor (cAMPi) H89 (10 mM) and a CREB inhibitor (CREBi) 666-15 (1 mM). MU41-CM with DMSO, and DMSO in fresh media (RPMI) ‘medium/med’, are negative controls, basal medium without GBM-CM.

Next, we investigated the potential link between CREB activation and upstream cell signaling pathways. To determine whether the cAMP-CREB axis is active in the GBM TME, we first performed multiplex immunohistochemistry using phospho-specific antibodies to identify PI3K and MAPK cell signaling activity in pCREB^+^ CD68^+^ TAMs, given the previously reported roles of these pathways in regulating macrophage polarization in the tumor microenvironment and in inflammation ^20,21^ (Fig 4D). Most TAMs in the GBM tumor microenvironment did not express pAKT and pERK1/2 (Fig 4E).

To test the potential role of upstream cell signaling pathways in CREB activation, in TAMs, we used a panel of pathway-specific small molecule inhibitors to target each pathway in GBM MU41-stimulated THP-1 macrophages, since conditioned medium from these patient-derived GBM cells exhibited robust CREB activation in macrophages (Fig 4A). Only the PKA inhibitor, H89, which suppresses activation of cAMP signaling, was able to inhibit CREB phosphorylation in GBM-stimulated THP-1 macrophages (Fig 4F). The inhibitors targeting the PI3K (BKM-120), MAPK (U0126) pathways and CREB-CBP interaction (666-15) had no effect on CREB phosphorylation in THP-1 macrophages. Collectively, the spatial tissue analysis and in vitro data show that the cAMP-CREB signaling-transcription axis regulates bone marrow-derived TAM immunosuppressive polarization, in response to factors secreted by GBM cells.

### GBM-secreted low molecular weight polypeptides regulate macrophage polarization

To determine the molecular properties of the GBM-secreted factors which regulate macrophage polarization in GBM, we performed targeted proteomic analysis of the MU41 GBM cell conditioned medium. First, using size exclusion filtration, we tested several molecular size ranges and showed that the most robust pCREB expression in THP-1 macrophages was mediated by factors smaller than 10kDa (Fig 5A). Moreover, proteinase K-treated unfiltered MU41 GBM cell conditioned medium had no effect on pCREB expression, nor CREB-dependent transcription (Fig 5B), demonstrating that the GBM-secreted factors able to regulate CREB transactivation in macrophages, are low molecular weight proteins, ranging between 10 and 30kDa. To identify proteins secreted by GBM cells which regulate macrophage polarization, we performed targeted Olink proteomic analysis using an Olink Target 96 Inflammation panel. Differential protein expression analysis was performed on GBM cell conditioned media produced by MU41 cell and U87 cells, to determine proteins expressed and secreted by GBM cell lines exhibiting either high (MU41) or low (U87MG) CREB activation in macrophages (Fig 5C; also refer to Fig 4A). Cytokine bead array assays confirmed the Olink proteomic analysis data, showing that MU41 cells expressed and secreted higher levels of several cytokines and chemokines, compared to U87MG cells, including C-C motif chemokine ligand (CCL20), C-X-C motif chemokine ligand 1 (CXCL1), monocyte chemoattractant protein-1 (MCP-1/CCL2) and C-C motif chemokine ligand 4 (CCL4) (Fig 5D). U87MG cells expressed and secreted higher levels some mitogenic and morphogenetic factors, including hepatocyte growth factor (HGF) and fibroblast growth factor 5 (FGF5); factors which distinguished U87MG cells from MU41 cells.

**Figure 5.**
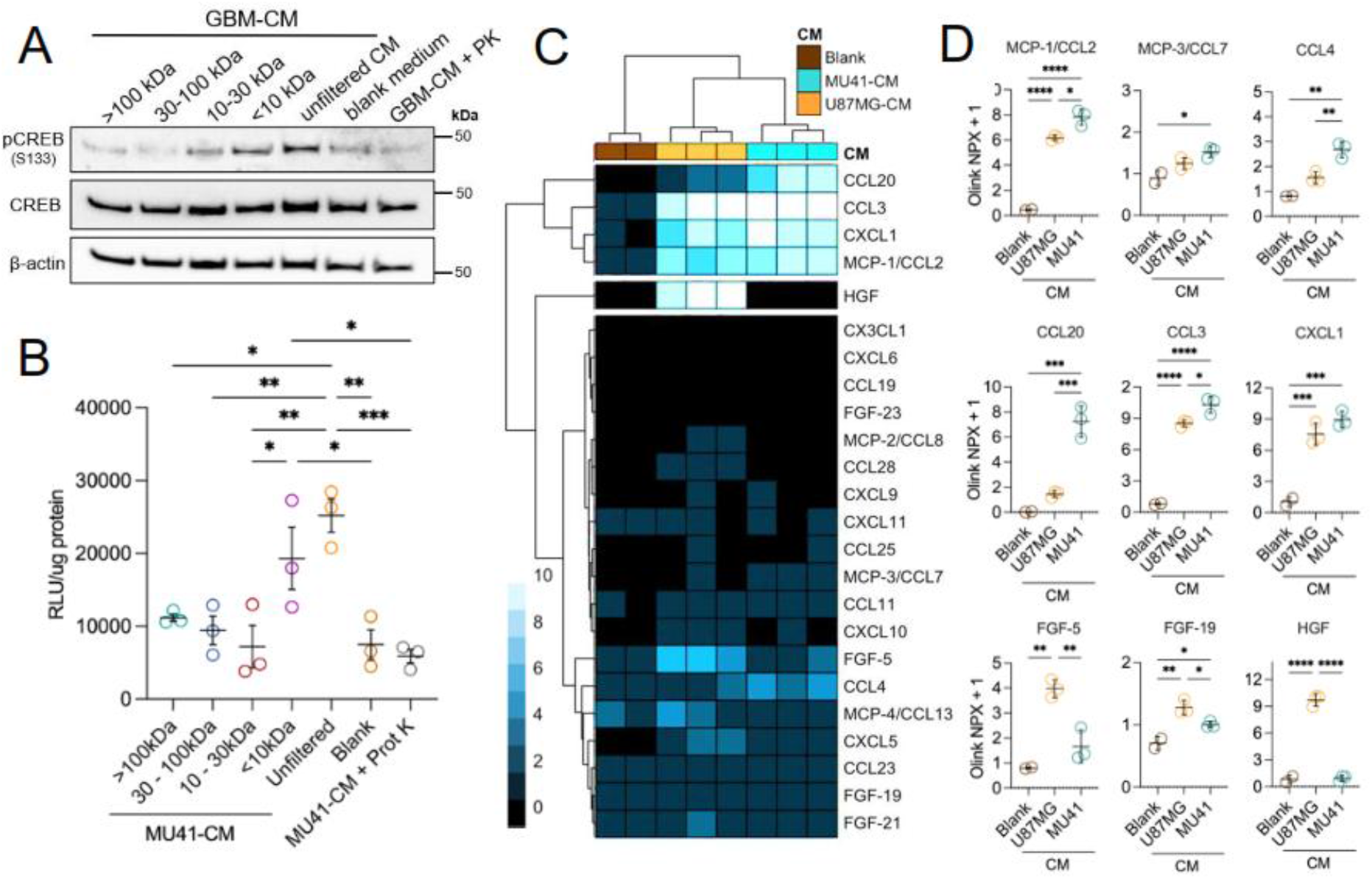
GBM-secreted macrophage modulating factors are low molecular weight proteins. **(A)** Cell lysates from THP-1 macrophages were probed for pCREB, total CREB (tCREB) and β-actin following a 30-minute exposure to size-based fractionated factors in MU41-CM. Unfiltered MU41-CM, fresh media (blank) and Proteinase K (PK)-treated MU41 GBM-CM lysate fractions were used as controls. **(B)** Experiment using fractionated MU41-CM was repeated on THP-1 macrophages harbouring a CRE/CREB reporter plasmid. Cell lysates were collected following a 7-hour exposure of the cells to fractionated CM and control medium. CREB activity is shown as RLU per mg protein. **(C, D)** The levels of chemokines and growth factors in MU41-CM, U87MG-CM, and a blank control without CM, were analysed using Olink Proteomics and presented as a heatmap, as Olink Normalised Protein eXpression (Olink NPX). Statistical significance was determined using one-way ANOVA, followed by Tukey’s test for multiple comparisons (a=0.05), and is represented by *(p<0.05), **(p<0.01), *** (p<0.001) and ****(p<0.0001). Error bars are median ± interquartile range.

### CREB target genes regulating GBM-stimulated macrophage polarization

To identify CREB-dependent target genes in bone marrow-derived TAMs which regulate immunosuppressive polarization we conducted experiments in GBM cell-stimulated macrophages generated from PBMC from six individuals (3 females and 3 males). PBMC macrophages were incubated with MU41 conditioned medium, with or without the CREB small molecule inhibitor, 666-15. RNA sequencing analysis was performed on RNA extracted from individual PBMC-macrophage cell preparations and Olink proteomic analysis was also conducted on conditioned media from these cells. For RNA sequencing analysis PBMC macrophages were incubated in the presence of MU41 conditioned medium, with or without CREB inhibitor 666-15, for 8 h, total RNA was then extracted and sequenced using the Illumina NovaSeq X Plus system, at a depth of greater than 20 million reads. For Olink proteomic analysis cells were stimulated with MU41 conditioned medium for 24 h, to account for the time difference required for RNA synthesis or protein synthesis, and to maximize the opportunity to measure mRNA and proteins encoded by CREB target genes (Fig 6A).

**Figure 6.**
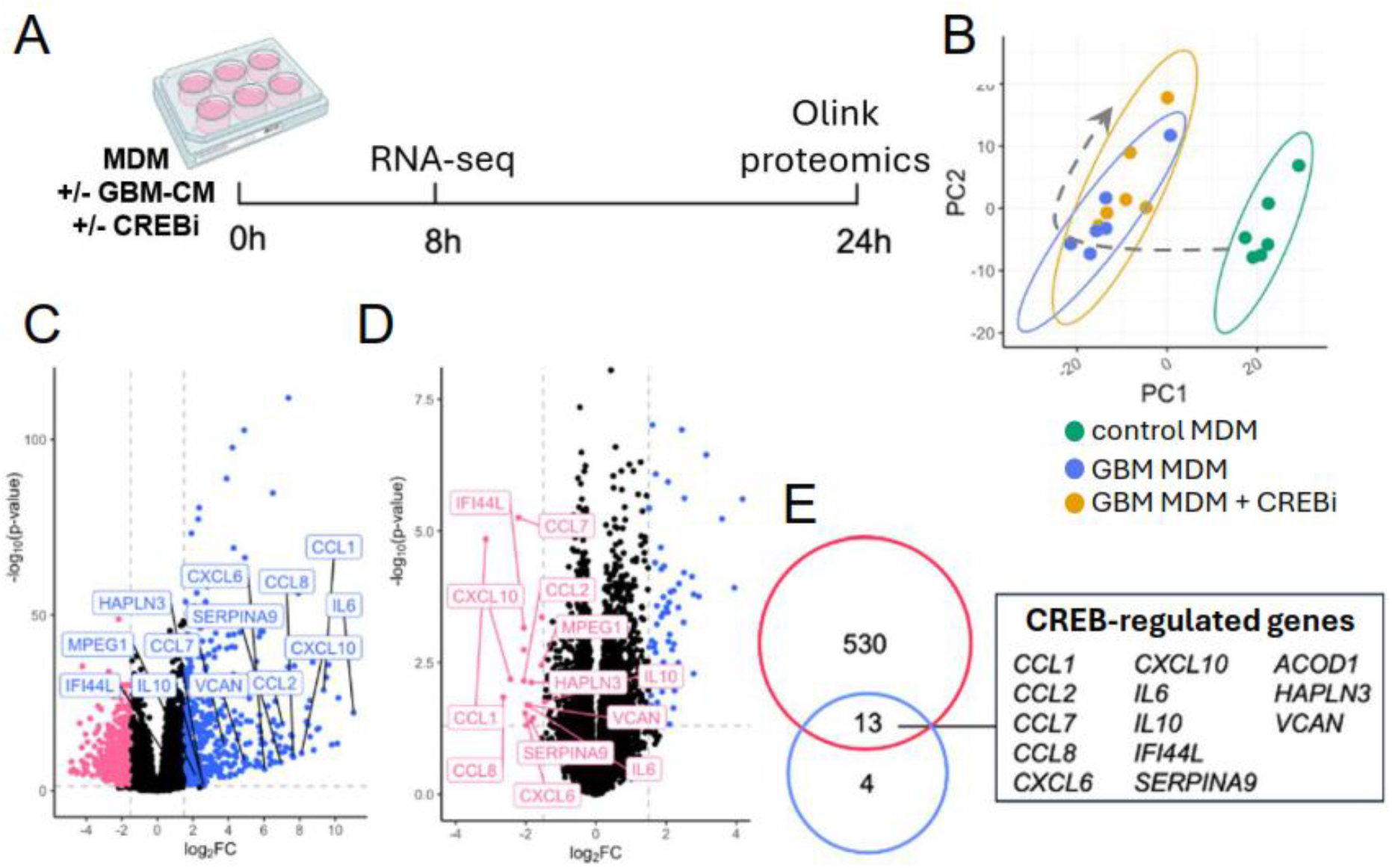
CREB regulates mRNA expression of anti-inflammatory factors and monocyte chemoattractant proteins in GBM TAMs. **(A)** CREB target gene analysis was performed by examining bulk RNA profiles of GBM TAMs treated with and without CREBi, and control macrophages, at 8 hours after GBM-CM and CREBi treatments. Secreted factors and surface biomarkers were also interrogated 24 hours following treatments using the Olink Target platform and flow cytometry, respectively. Diagram was created in Biorender. **(B)** Total RNA profiles of control macrophages (green), GBM-TAMs (light blue) and CREBi-treated GBM-TAMs (orange) are visualised in a principal component analysis (PCA) plot. **(C)** Volcano plots displaying upregulated and downregulated genes between GBM TAMs and control macrophages. **(D)** GBM TAMs with and without the CREBi. Dashed lines indicate the cutoff P-value and fold-change (FC) scores. **(E)** mRNA transcripts upregulated in MDM upon exposure to GBM-CM (GBM-TAM) (blue circle); transcripts downregulated in GBM-TAM treated with 1mM CREB inhibitor, 666-15 (red circle); the 13 transcripts common to both groups, are those which are CREB-regulated genes, listed in the text box.

Principal component analysis shows that the six individual PBMC macrophages can be separated based on their transcriptomic profile, where control untreated macrophages form a discrete non-overlapping cluster (Fig 6B), separated from GBM stimulated macrophages (GBM TAM), with or without CREB inhibitor. Differentially expressed genes were visualized using volcano plots to identify differences between control unstimulated macrophages and MU41 conditioned medium stimulated macrophages (Fig 6C), and between MU41 stimulated cells, with or without incubation with the CREB inhibitor (Fig 6D). PBMC macrophages stimulated with MU41 GBM cell conditioned medium exhibited significant (p>0.05), greater than 2-fold (log_2_FC >/=1) upregulation of 530 genes, compared to control, unstimulated cells.

We identified 17 transcripts expressed by MU41-stimulated GBM TAM which were downregulated in the presence of the CREB inhibitor, 666-15 (Fig 6D). The mRNAs encode for immunomodulatory factors including *IL10, CXCL10, CCL2, CCL8* and *CCL6*, the inflammatory inhibiting protein aconitate decarboxylase 1 (*ACOD1*), the macrophage survival/anti-apoptotic factor Serpin Family A Member 9 (*SERPINA9*), as well as the extracellular matrix proteins, versican (*VCAN*) and hyaluronan and proteoglycan link protein 3 (*HAPLN3*). The differentially expressed gene data is summarized in Fig 6E, which shows that 13 mRNAs expressed in macrophages are upregulated by GBM cell conditioned medium and downregulated by CREB inhibition, defining these as likely CREB target genes.

To determine whether GBM TAM mRNAs, identified by RNA-sequencing are translated and secreted, we performed protein analysis on GBM-macrophage conditioned medium using an Olink Target 96 Inflammation panel. Principal component analysis showed that GBM TAMs are separated from control untreated macrophages based on their secreted protein profiles and treatment with the small molecule CREB inhibitor shifts the profiles of these macrophages (Fig 7A).

**Figure 7.**
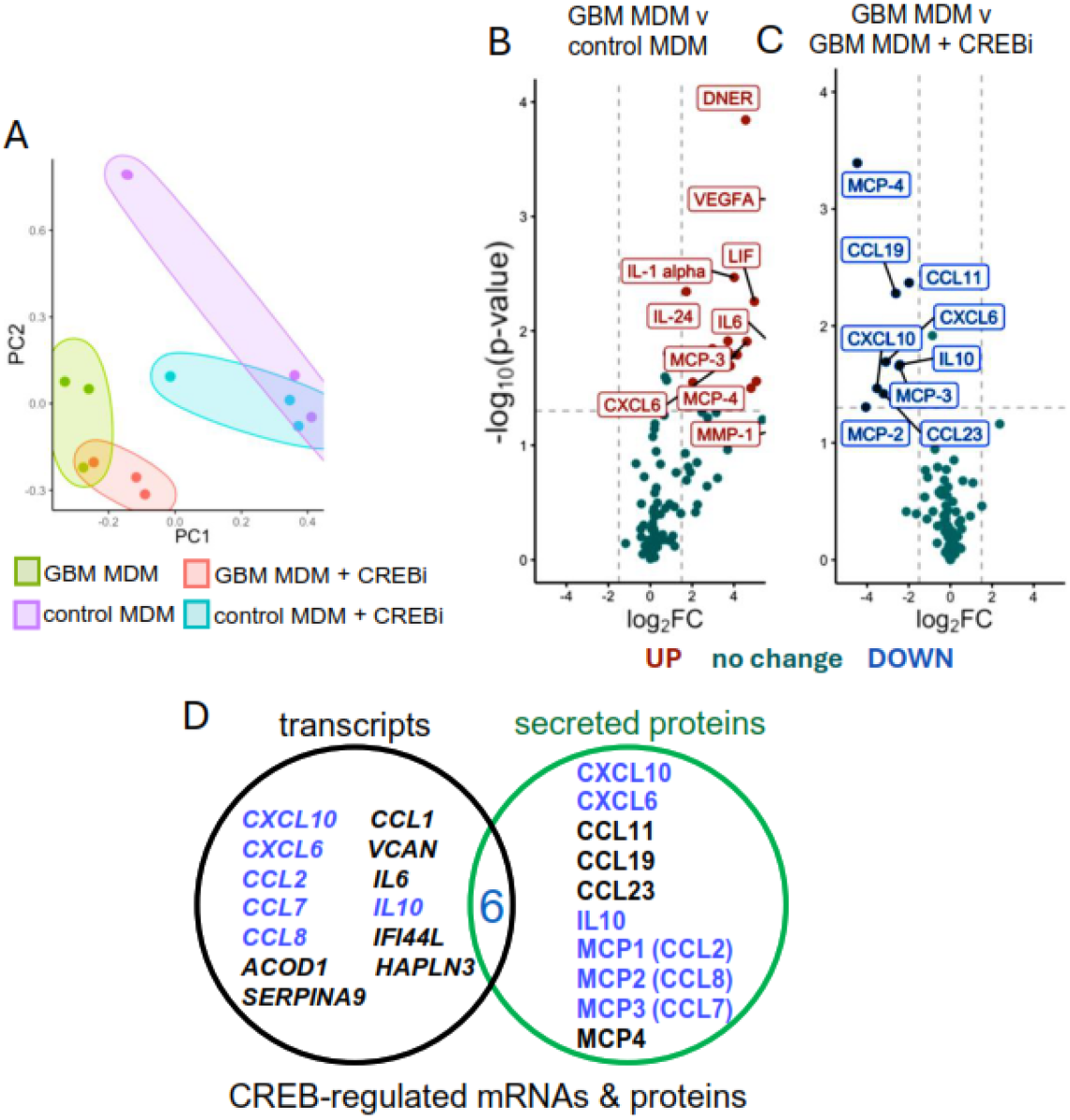
CREB regulates the expression of secreted immunosuppressive chemokines in GBM-CM stimulated macrophages. **(A)** Secreted protein data (Olink Target 96 Inflammation panel) principal component analysis (PCA) plot, for control macrophages (mΦ) and GBM-stimulated macrophages (GBM TAM), incubated with and without CREB inhibitor (CREBi). Experiments were preformed using four biological replicates (primary macrophages from four healthy donors). **(B)** Volcano plots show the expression differences of differentially secreted factors, between control control PBMC-macrophages and GBM-stimulated macrophages (GBM TAM), with or without CREBi. The dashed lines indicate significantly upregulated proteins p=0.05; log2FC=1.75 **(C)** CREB target genes and secreted proteins in GBM-stimulated macrophages showing the transcripts identified by RNAseq and Olink proteomic analysis. The CREB-regulated factors identified by both RNAseq analysis and Olink proteomic analysis are shown in blue font. **(D)** Proteins regulated by CREB. A schematic diagram showing CREB-regulated expression of 5 proteins identified using Olink Target 96 Inflammation panel protein analysis and 4 proteins identified by cytometric-bead array analysis, defined by their upregulation in GBM TAMs relative to control macrophages, and downregulation in GBM TAMs treated with CREBi, compared to GBM TAMs without CREBi.

Differential protein expression analysis shows the secreted cytokines and chemokines which are upregulated in macrophages in response to GBM conditioned medium stimulation. The secreted proteins which exhibit significant upregulation include MMP1, CXCL6, MCP-4, MCP-3, IL6, IL24, IL1α, LIF, VEGFA and DNER (Fig 7B). A comparison of the differential protein expression and secretion, between GBM-stimulated macrophages incubated with and without the CREB inhibitor, 666-15, showed that nine cytokines and chemokines were downregulated in response to CREB inhibition: CCL23, MCP-2, MCP-3, IL10, CXCL10, CXCL6, CCL19, CCL11, MCP-4 (Fig 7C). To confirm the Olink analysis of secreted factors, cytometric bead arrays (CBA) were performed for a panel of cytokines (Fig S1). Of the secreted proteins, six were also identified by RNA-sequencing: CXCL10, CXCL6, IL-10, CCL2, CCL8 and CCL7 (Fig 7D).

## Discussion

Suppression of anti-tumor immunity has implications in immunotherapy, where the best outcomes for patients with GBM across multiple clinical trials investigating various immunotherapeutic strategies have led to transient responses, restricted to very few patients^22^. This lack of success can be attributed to multiple factors, with macrophage-mediated immunosuppression considered to be prominent, given the abundance and efficient infiltration of MDM into the tumor parenchyma^2^. TAMs exist in a state of flux, transitioning between ‘anti-tumor’ inflammatory states, and ‘pro-tumor’ immunosuppressive states. Thus, there is a major research effort focused on reprogramming macrophages in GBM. However, the mechanisms regulating macrophage biology in the GBM TME, including understanding how TAMs become immunosuppressive and maintain this phenotype throughout the development of tumors, remain to be elucidated. In this study, we aimed to define factors promoting the transition of macrophages toward immunosuppression in GBM and to identify cell signaling pathways and transcription factors which regulate this transition.

Several studies have reported that the perivascular niche in GBM is rich in immunosuppressive macrophages, lymphocytes and ECM proteins^18,23,24^. Using spatial analysis, we observed that although both bone marrow-derived TAMs and lymphocytes are present in GBM tumors, only TAMs exhibit efficient infiltration beyond the perivascular niche and into the tumor parenchyma, compared to lymphocytes. Whether or not there is a dynamic shift in macrophage polarization between the circulating macrophages and those in the GBM tumor parenchyma in patients with GBM, has thus far, not been resolved. Spatial infiltration analysis to compare macrophage polarization status in tumor vessels, and within and beyond perivascular niches, showed that bone marrow-derived TAMs arrive as inflammatory cells and as they enter the tumor parenchyma, they polarize toward an immunosuppressive state. A study investigating the TME in PDGF-driven and GL261 GBM mouse models showed that monocytes arriving via the circulation are inflammatory.

However, this study did not determine the polarization status of macrophages within the brain tumor parenchyma^24^.

To determine bone marrow-derived TAMs activation status we measured changes in expression of the kinase inducible transcription factor CREB. Our previous work demonstrated that targeting CREB in a Pi3k-Pten GBM mouse model resulted in slower tumor growth and extended symptom free survival^25^. Here and in our previous study, we show that in GBM tumors, the proportion of both T-cells and bone marrow-derived TAMs positive for phosphorylated CREB (pCREB^+^) increases during disease progression, comparing lower grade astrocytoma to higher grade astrocytoma and GBM, irrespective of IDH mutation status (Dinevska et al., 2023). We also demonstrate a change in CREB activation in MDM, dependent on their localization. pCREB^+^ TAMs in the tumor parenchyma are immunosuppressive, whereas vascular and perivascular are pCREB negative. This observation suggests that once MDMs exit the blood vessels and encounter the perivascular tumor niche, CREB is activated in MDMs via upstream signaling leading to an increase in pCREB expression. Our data demonstrated that factors secreted by GBM cells activate CREB in MDMs and that these GBM derived factors are sufficient to stimulate TAM polarization toward an immunosuppressive phenotype. Previous studies have shown that glioma-secreted factors and exosomes are able to promote bone marrow-derived TAM immunosuppression^26,27^. Our study also showed that GBM-secreted factors able to regulate TAM immunosuppression are proteins smaller than 10kDa, and need not be associated with exosomes, since exosomes would be excluded in the macrophage-active 10kDa conditioned medium filtered fraction, used in our study. This conclusion is supported by several studies demonstrating that exosomes derived from various cultured cells range between 40 to 150 nm in diameter, and are effectively retained using protein filters with a molecular weight cut-off of 10kDa, thus excluded from resulting flow-through filtrates ^28,29^.

An important finding of our study was that pharmacological inhibition of CREB reverses TAM immunosuppression, in vitro. By examining several upstream cell-signaling pathways which potentially activate CREB in TAM, we showed that cAMP/PKA signaling is the pathway which activates CREB via phosphorylation, and that the PI3K and MAPK/ERK pathways are not prominently activated in GBM tumor tissue. Previous work has linked either the cAMP pathway or a cAMP-CREB axis with TAM immunosuppression in neuroinflammation or breast cancer ^15,30–32^. The PI3K/AKT pathway has also been shown to be activated by GBM secreted factors in mouse macrophages, in vitro^26^. Differences in cell signaling activation and TAM polarization in different studies highlight the complexity of GBM cell-macrophage communication, and differences in the factors secreted by GBM cell lines. In this study, GBM cell lines U87MG and T98G, which are poorly tumorigenic in vivo and exhibit low invasive capacity, secrete high levels of mitogenic factors but not immunomodulatory factors, with the opposite observed in highly invasive and tumorigenic GBM cell lines, LN229 and patient-derived MU41 cells.

Many intracellular cell-signaling and transcriptional mechanisms regulating TAM pro-tumor polarization have been examined. Hypoxia is a cancer hallmark of GBM and single cell sequencing studies show that the level of hypoxia correlates with increased suppression of both macrophages and T-cells in GBM, and that the transcription factor hypoxia-inducible factor-1α likely regulates the expression of immunosuppressive genes in TAMs in GBM ^33^. Single cell CRISPR screening identified the transcription repressor, ZEB2 as a master regulator of TAM immunosuppression across multiple cancer types, including GBM, and that targeting ZEB2 reprograms mouse TAMs to induce anti-tumor immunity^34^. In our study, we show that transcriptional activation mediated by CREB regulates TAM immunosuppressive polarization by activating the transcription of thirteen genes encoding secreted cytokines and chemokines. These include eleven immunomodulatory factors, IL10, CCL2 and CXCL10, and two ECM or ECM modifying proteins, VCAN and HAPLN3, respectively. Among these factors, IL10 expression has been extensively studied and shown to be regulated by cAMP and CREB via several well characterized CREB response element (CRE) sequences in the *IL10* promoter ^13,35^. Our previous studies have shown that extensive tumor ECM remodeling occurs in GBM and metastatic brain tumors, and that the remodeling alters tumor immune cell localization and function, so combined with secretion of immunomodulatory factors, brain tumor cells exploit macrophage functions to thrive^18,36^. Further, the data in this study shows that macrophage cell surface proteins regulated by CREB also define pro-tumor immunosuppressive functions.

Previous work targeting CREB in human myeloid leukemia cells using shRNA and microarray analysis has identified CREB target genes^37^. To our knowledge, no other studies have systematically investigated CREB target genes in TAMs, nor primary monocytes/macrophages. Comparison of CREB target genes identified by Pellegrini et al., and other studies using other cell types, with the CREB target genes identified in TAMs, we show that there are common CREB target genes across multiple studies (Table S2).

Overall, our study defines key GBM secreted immunomodulatory factors which promote macrophage immunosuppression via the cAMP-CREB signaling-transcription axis and provides a mechanistic framework for targeting neoplastic cells and reprogramming the tumor microenvironment to enhance anti-tumor effects.

## Acknowledgments

We thank Professor Christine Wells for ongoing scientific advice. We thank Associate Professor Nichollas E. Scott for assistance with the proteomic analysis. We acknowledge the use of the services and facilities of AGRF and the University of Melbourne Cytometry Platform (Doherty Institute and Melbourne Brain Centre nodes) for provision of flow cytometry services. This work was supported by the following grants: SSW was supported by a University of Melbourne, Melbourne Research Scholarship. MD was supported by an Australian Government Research Training Program Scholarship. MH, LS & TM were supported by an NHMRC Ideas grant (2021360). TM was supported by a CASS Foundation Science Grant (065516), a Brain Foundation Grant (065518) and a Department of Microbiology & Immunology Collaborative Grant, and the Department of Surgery (RMH), The University of Melbourne. SSS was supported by an MRFF Accelerated Research-Australian Brain Cancer Mission grant (1158175). LC was supported by an NHMRC Investigator Grant (2009339) and a Department of Microbiology & Immunology Collaborative Grant, The University of Melbourne.

## Author contributions

SSW, MD, SSS, LC and TM conceived and supervised the study. and drafted the manuscript. TM funded the study. SSW and MD also performed the spatial analysis and prepared the figures. RD and RM assisted with cell biology, flow cytometry experiments and finalization of the manuscript. PF and TCCL performed the Olink proteomic experiments. SM assisted with the RNA sequencing analysis. LC and LAA assisted with the primary monocyte cell biology and flow sorting experiments. SV and JT assisted with data analysis and manuscript writing. MH and LS co-funded parts of the study and assisted with the data analysis and manuscript writing. MABR and AB provided key reagents and assisted with cell biology experiments and manuscript writing.

## Supplementary Data

**Table S1:** Antibodies for multiplex immunohistochemistry *CST - Cell Signaling Technology.

| Antibody | Source | Catalogue # | Dilution |
| --- | --- | --- | --- |
| anti-CD3 [SP7] | Abcam | CD16669 | 1:100 |
| anti-CD68 [KP1] | Abcam | ab955 | 1:100 |
| anti-TMEM119 – C-terminal | Abcam | ab185333 | 1:500 |
| CD31 JC70A | Dako | M0823 | 1:40 |
| HLA-DRA (E9R2Q) XP | CST* | #97971 | 1:250 |
| CD163 (D6U1J) | CST | #93498 | 1:200 |
| Anti-Mannose Receptor | Abcam | ab64693 | 1:2000 |
| Recombinant anti-pCREB<br>(S133) [E133] | Abcam | ab32096 | 1:1000 |
| phospho-Akt (Ser473) (D9E) XP | CST | #4060 | 1:100 |
| phospho-p44/42 MAPK (Erk 1/2)<br>(Thr202/Tyr204) (D13.14.4E)<br>XP | CST | #4370 | 1:200 |

\*CST - Cell Signaling Technology

**Table S2:** CREB target genes with experimentally validated or predicted cAMP Response Element (CRE) promoter binding sites.

| **CREB Target Genes** | **Method** | **Reference** |
| --- | --- | --- |
| IL10 | promoter point mutations and CRE-luciferase reporter activity | 13 |
| CXCL10, CXCL6, CCL7, CCL11, IL6 | shRNA CREB knockdown and microarray analysis | 37 |
| ACOD1, HAPNL3, IFI44L, DNER | GeneHancer - Predicted | 38 |
| DNER | ECODE Project - Predicted | 39 |

## Supplementary data

**Figure S1.**
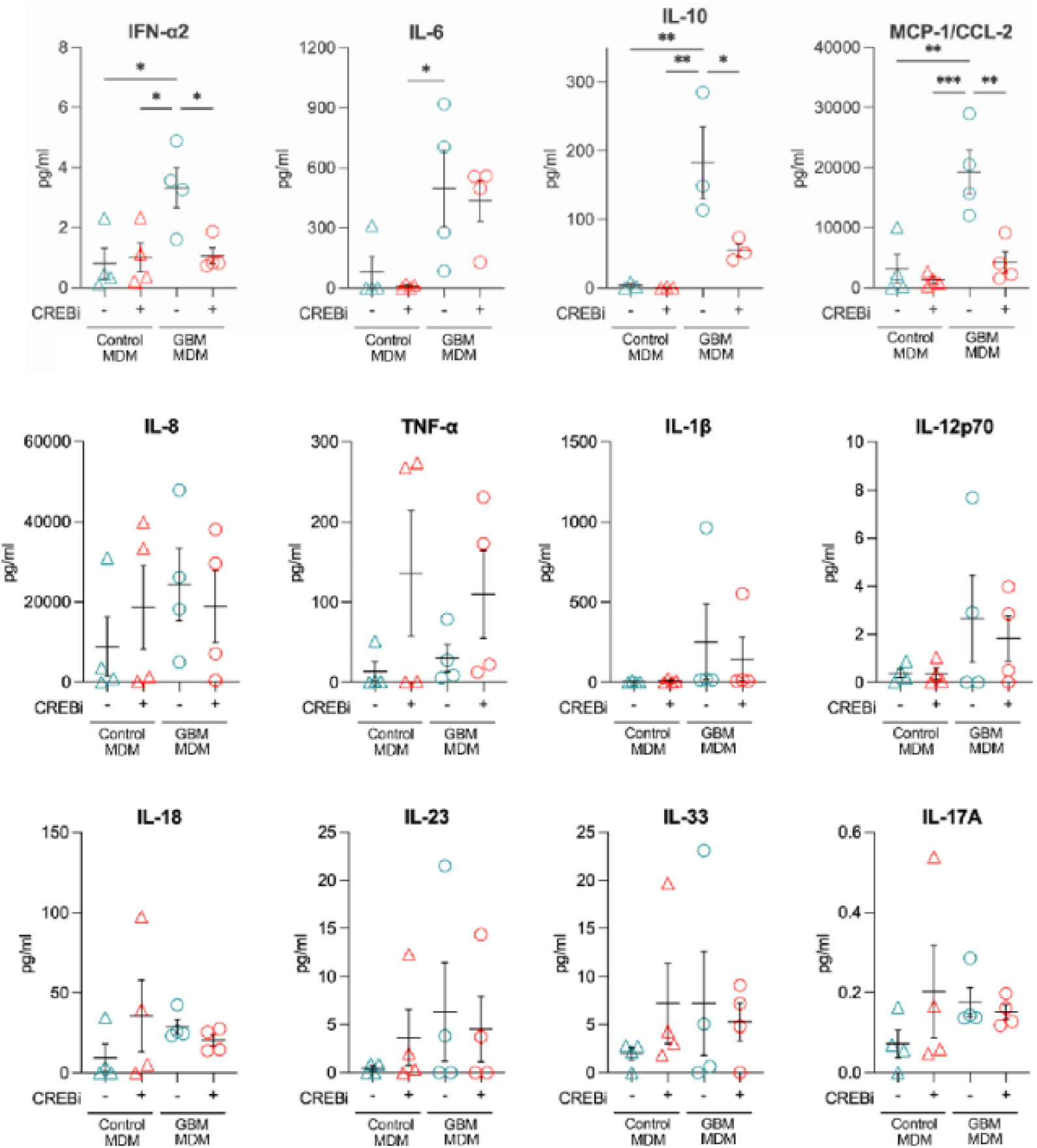
CREB-regulated secreted cytokines and chemokines in GBM TAMs. Cytokines and chemokines secreted by GBM TAMs and control macrophages with/without CREBi, measured using a cytometric bead array, 24h post-treatment. Experiments were performed on four primary macrophage preparations from 4 healthy blood donors. Statistical significance was determined by an ordinary one-way ANOVA and is represented by *(p<0.05), **(p<0.01), *** (p<0.001) and ****(p<0.0001).

